# Morphology and connectivity of retinal horizontal cells in two avian species

**DOI:** 10.1101/2025.01.27.634460

**Authors:** Anja Günther, Vaishnavi Balaji, Bo Leberecht, Julia J. Forst, Alexander Y. Rotov, Tobias Woldt, Dinora Abdulazhanova, Henrik Mouritsen, Karin Dedek

## Abstract

In the outer vertebrate retina, the visual signal is separated into intensity and wavelength information. In birds, seven types of photoreceptors (one rod, four single cones, and two members of the double cone) mediate signals to >20 types of second-order neurons, the bipolar cells and horizontal cells. Horizontal cells contribute to color and contrast processing by providing feedback signals to photoreceptors and feedforward signals to bipolar cells. In fish, reptiles, and amphibians they either encode intensity or show color-opponent responses. Yet, for the bird retina, the number of horizontal cell types is not fully resolved and even more importantly, the synapses between photoreceptors and horizontal cells have never been quantified for any bird species. With a combination of light microscopy and serial EM reconstructions, we found four different types of horizontal cells in two distantly related species, the domestic chicken and the European robin. In agreement with some earlier studies, we confirmed two highly abundant cell types (H1, H2) and two rare cell types (H3, H4), of which H1 is an axon-bearing cell, whereas H2-H4 are axonless. Horizontal cell contacts to photoreceptors were type-specific and similar to the turtle retina, which confirms the high degree of evolutionary conservation in the vertebrate outer retina. Our data further suggests that H1 and potentially H2 cells may encode intensity, whereas H3 and H4 may represent color opponent horizontal cells which may contribute to the birds’ superb color and/or high acuity vision.

## 1 Introduction

The vertebrate retina contains five classes of neurons: photoreceptors transduce the incoming visual signals and propagate them via bipolar cells to retinal ganglion cells. Horizontal cells modulate the signals from photoreceptors to bipolar cells in the outer retina whereas amacrine cells modify signals between bipolar cells and ganglion cells in the inner retina. To fulfill the complex functions of the retina, for example luminance, color, contrast, and motion processing (Kerschensteiner, 2022) and, in at least some bird species, light-dependent magnetoreception (Chetverikova et al., 2022; Günther et al., 2018; Xu et al., 2021; Zapka et al., 2009), each of the five classes of neurons comprises many different cell types (Hahn et al., 2023; with few species-specific exceptions). In the mouse retina, the various cell types have been distinguished based on morphology (Sümbül et al., 2014; Wässle et al., 2009), connectivity (Behrens et al., 2016; Helmstaedter et al., 2013), gene expression (Macosko et al., 2015; Shekhar and Sanes, 2021), and physiology (Baden et al., 2016; Hsiang et al., 2024).

However, much less is known in the bird retina, despite these recent advances: a transcriptomics-based cell atlas revealed 133 different neuronal cell types in the chicken retina (Yamagata et al., 2021), serial sectioning volume electron microscopy reconstructions of the chicken and European robin retina showed a highly complex connectivity between photoreceptors and unique sets of bipolar cell types (Günther et al., 2024, 2021), a screen for immunohistological markers revealed pronounced variation in marker signature for the bipolar cells of different bird species (Balaji et al., 2023), and the first systematic overview of retinal ganglion cell responses from the chicken retina demonstrated striking differences in signal processing between mammalian and avian retinas (Seifert et al., 2023). Yet, detailed information on their connectivity is still missing for many cell types in the bird retina.

Here, we focus on the horizontal cells and their connections with photoreceptors. We studied two distantly related bird species, the domestic chicken (*Gallus gallus domesticus*) and the night-migratory European robin (*Erithacus rubecula*) which inhabit different ecological niches and forage differently. Furthermore, the European robin is known to have a light-dependent magnetic compass located in the retina (Chetverikova et al., 2022; Günther et al., 2018; Wiltschko et al., 1993; Xu et al., 2021; Zapka et al., 2009). Horizontal cells are essential to establish and maintain the photoreceptor/bipolar cell synapse (Nemitz et al., 2021, 2019; Sonntag et al., 2012). They also play an important role in retinal signal processing (Chaya et al., 2017; Drinnenberg et al., 2018; Ströh et al., 2018) by providing feedback signals to photoreceptors (reviewed in Thoreson and Mangel, 2012) and feedforward signals to bipolar cells (Behrens et al., 2021). Their potential function in light-dependent magnetoreception is completely unknown. Horizontal cells can be divided into axon-bearing and axon-less cell types, a division, which appears to be conserved among vertebrates (Boije et al., 2016; Hahn et al., 2023; Peichl and Gonzalez-Soriano, 1994). In teleost (e.g., zebrafish: Connaughton and Nelson, 2010; Yoshimatsu et al., 2021), reptiles (turtles: Ammermüller et al., 1995; Asi and Perlman, 1998), and amphibians (frog: Witkovsky, 2000) some horizontal cell types respond with different polarity to spectral stimuli (color-opponent or chromaticity horizontal cells), whereas others encode changes in brightness, irrespective of stimulus color (luminosity horizontal cells) (reviewed in Thoreson and Dacey, 2019). Whether a similar division is present in the bird retina is unclear so far. In the chicken retina, the number of horizontal cell types is not fully resolved yet (reviewed in Seifert et al., 2020). While one type of axon-bearing horizontal cell has been confirmed (termed H1), two axon-less types have been described based on Golgi impregnation (H2-3; Gallego, 1986) and membrane-targeted EGFP expression after *in-ovo* electroporation (Tanabe et al., 2006). However, based on immunohistochemical staining, three different types of axon-less horizontal cells were reported (H2-4; Fischer et al., 2007), in line with the recent chicken transcriptome dataset (Yamagata et al., 2021).

To resolve these discrepancies, we immunostained the chicken retina following Fischer et al. (2007) and confirmed the existence of four different types of horizontal cells (H1-H4) across the entire tissue. However, using the same markers we were only able to distinguish three different types in the European robin retina. Therefore, we used a previously published volume electron microscopy dataset to reconstruct the horizontal cells in the dorsal periphery (Günther et al., 2024). We 1) identified four types of horizontal cells in the dorsal European robin retina, 2) revealed their distinct connections to the different photoreceptor types to generate predictions on their physiological responses, and 3) found unusual chemical synapses (mainly in H1 horizontal cells), which target distinct bipolar cell types and an interplexiform amacrine cell.

## 2 Materials and methods

### 2.1 Animals and tissue preparation

Domestic chickens were obtained as 1-day-old chicklings from a local breeder (Brüterei-Siemers GmbH & Co. KG, Lohne, Germany) and raised in the animal facility of University of Oldenburg (Oldenburg, Germany). Adult European robins were caught using mist nets in the vicinity of University of Oldenburg; catching was performed based on a permit from the Lower Saxony State Department for Waterway, Coastal and Nature Conservation (D7.2220/18). Birds were housed indoors under the natural light-dark cycle in the animal facility of University of Oldenburg with *ad libitum* access to food and water. If under 250 g, chickens (aged >2-weeks) and adult robins were sacrificed by decapitation; heavier chickens were euthanized by an overdose of Narcoren or Narkodorm before decapitation. All animal procedures were performed in accordance with local, national and EU guidelines for the use of animals in research and were approved by the Animal Care and Use Committees of the *Niedersächsisches Landesamt für Verbraucherschutz und Lebensmittelsicherheit* (LAVES, Oldenburg, Germany).

After sacrifice, eyes were quickly removed from the head and placed into a petri dish containing 30-35 °C carboxygenated Ames’ medium (A1420-10X1L, Sigma-Aldrich, St Louis, USA) supplemented with additional 30 mM NaHCO_3_, 25 mM glucose, 2.4 mM KCl and 1.8 mM MgCl_2_ or extracellular solution (in mM: 100 NaCl, 6 KCl, 1 CaCl_2_, 2 MgSO_4_, 1 NaH_2_PO_4_, 30 NaHCO_3_, 50 glucose, pH 7.4). The anterior part of the eye was cut with a razor blade and separated from the rest of the eyecup; the vitreous was carefully removed.

### 2.2 Fluorescent dye injection in vibratome slices of the chicken retina

For intracellular injections of horizontal cells, the chicken retina was gently dissected from the eyecup, separated from the pigment epithelium in extracellular solution and cut into small pieces which were embedded in low-melting point agarose (1.5% in extracellular solution, Bio & Sell GmbH, Feucht, Germany, #BS20.47.025). Vertical sections (200 µm) were cut with a vibratome (Leica VT 1200 S, Leica Biosystems, Wetzlar, Germany) and briefly fixed in 2% paraformaldehyde (PFA, in phosphate buffered saline, PBS) for 6.5 minutes. After washing, the slices were placed in a bath chamber and the horizontal cell layer was identified using a 40× or 63× water immersion objective under bright light illumination. For fluorescent dye injection, electrodes were made from borosilicate capillaries (Hilgenberg, Malsfeld, Germany) with a P-97 electrode puller (Sutter Instruments) and had resistances between 120-180 MΩ. They were tip-filled with 2 µl of Alexa Flour 488 hydrazide or Alexa Flour 568 hydrazide (dissolved in 200 mM KCl) and backfilled with 8-10 µl 200 mM KCl. Horizontal cells were targeted under visual control and impaled for dye iontophoresis as described earlier (Tetenborg et al., 2017), using a Multiclamp 700B amplifier (Molecular Devices) generating 500 ms long square pulses with 2 nA at 1 Hz for 2 min. After injection, the dye was allowed to diffuse for at least 15 min before the slice was fixed in 2-4% PFA for 15 min and washed in PBS.

### 2.3 Immunohistochemistry

For immunohistochemistry of vertical retina sections, the eyecups were immediately immersion-fixed in 4% PFA in PBS for 30 min. Afterwards, they were washed in PBS and subsequently passed through 10%, 20% and 30% sucrose solutions in PBS until the eyecups sank. Vertical sections (25 µm) were cut on a cryostat, dried on a hot plate for >45 min and stored at -20° C until further use. Slices were washed three times for 10 minutes in PBS and incubated in blocking solution containing 3% donkey serum and 0.3% Triton-X in PBS for 1 h. Primary antibodies (Table 1) were applied in blocking solution overnight at 4° C. Slices were washed in PBS and subsequently incubated with secondary antibodies (Table 1) diluted in the blocking solution for 2 h at room temperature. After washing, slices were mounted in a few drops of Vectashield antifade medium containing 4′,6-diamidino-2-phenylindole (DAPI, H-1200-10, Vector Labs) and covered with #1.5 glass coverslips. In control stainings, we omitted the primary antibodies to test for unspecific binding of the secondary antibodies. No unspecific staining was detected.

**Table 1.**
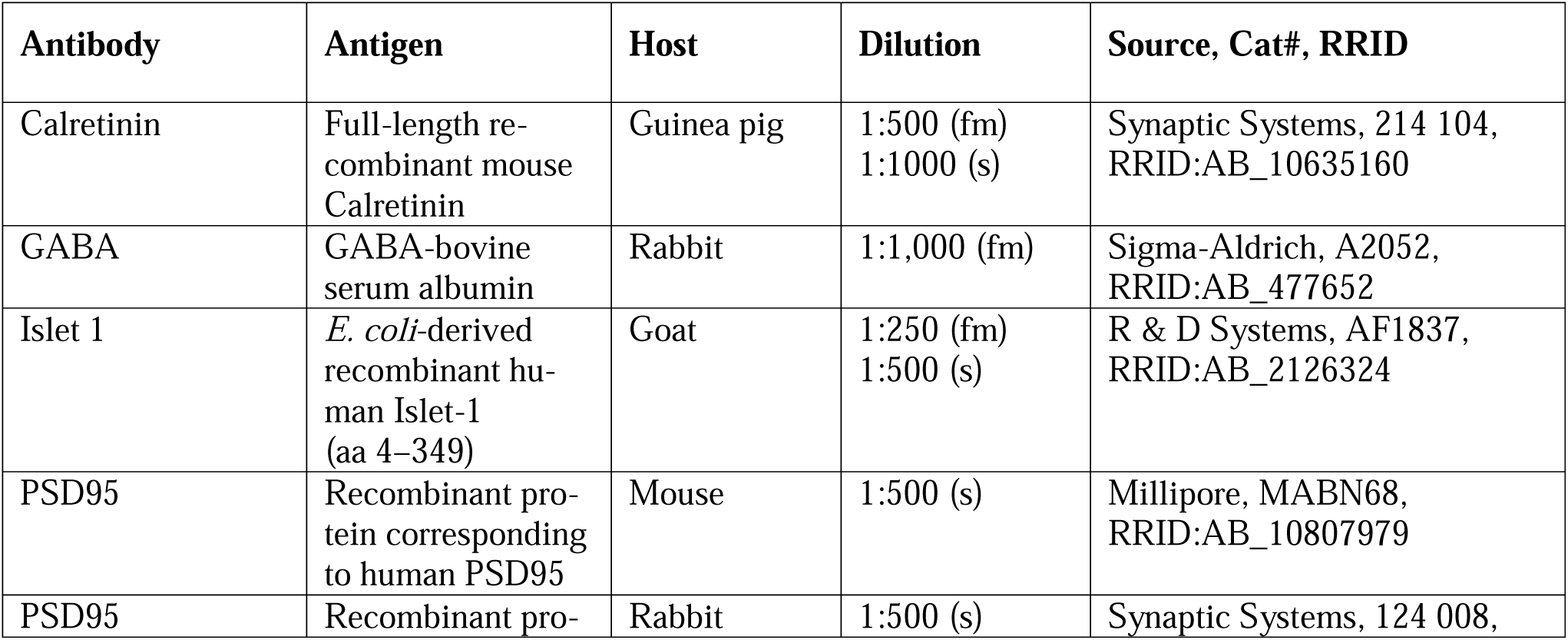
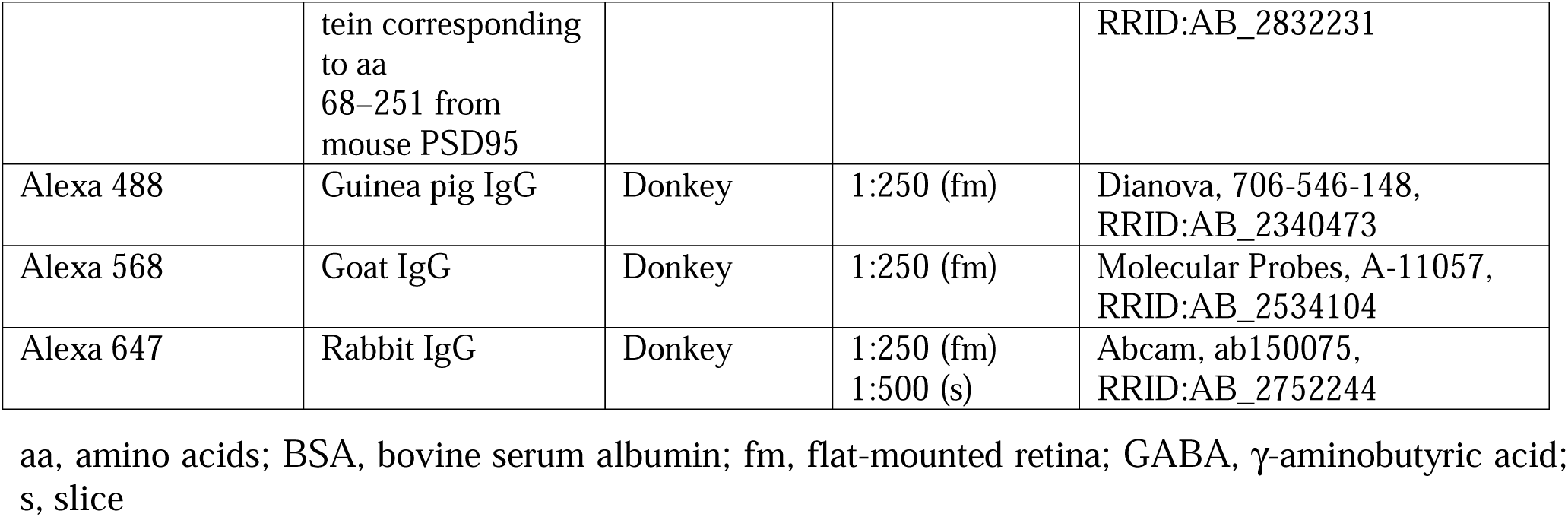
Primary and secondary antibodies used in this study.

For immunohistochemistry of flat-mount retinas, the pecten was dissected away before the retina was removed from the eyecup. For proper flat-mounting, 2-3 small incisions were made in the dorsalventral and nasal-temporal periphery regions (Fig. 1B). Then, the retina was flat-mounted photoreceptor side up on black filter paper (AAWP04700, MF-Millipore, Merck, Germany). The flat-mounts were fixed in 4% PFA in PBS for 30 min. After two washes in PBS for 30 min each, the flat-mounts were immersed in PBS containing 10%, 20%, and 30% sucrose for 30 min each. The flat mounts were frozen in PBS containing 30% sucrose and stored at -20°C until further use. To better visualize horizontal cells in flat-mounted tissue, the retina flat-mounts were thawed and put into a petri dish with PBS. Under a stereomicroscope, the outer segments of photoreceptors were gently brushed away using a thin paintbrush until the first oil droplets were brushed away.

**Figure 1.**
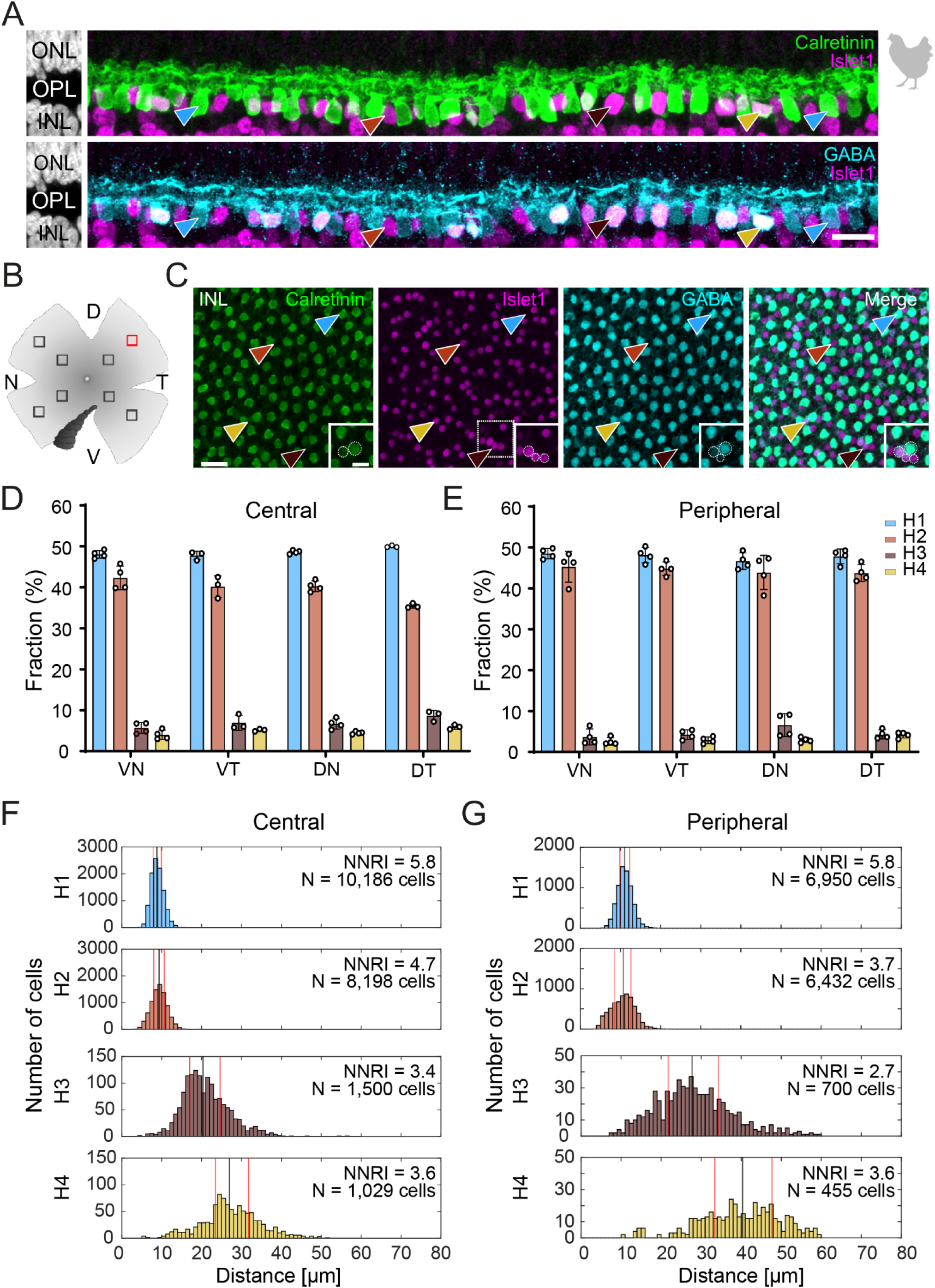
Triple labeling with Calretinin, GABA, and Islet1 allows the differentiation of four types of horizontal cells in the chicken retina. **(A)** Maximum projections of vertical slices of the chicken retina labeled for Calretinin (Calret), GABA, and Islet1. Based on these markers, four different types of horizontal cells can be distinguished: H1: Calret+/GABA+ (blue arrowhead); H2: Islet1+ (red arrowhead); H3: Islet+/GABA+ (brown arrowhead), and H4: Calret+/GABA+/Islet1+ (yellow arrowhead). **(B)** Schematic of the flat-mounted retina of a chicken with cuts in the nasal (N), dorsal (D), temporal (T) and ventral (V) side to flatten the tissue. The dark comb-like structure represents the pecten. Four image stacks each were taken from the central and peripheral retina, marked by the squares (not to scale). An example from the dorsal-temporal region (red square) is shown in (C). **(C)** Flat-mounted retina from the chicken retina, labeled for Calret, GABA and Islet1. Focus is on the INL. Arrowheads point to the different types of horizontal cells with the same color code as in (B). The area marked with a dashed square is shown enlarged in the insets for all three channels; this area contains all four horizontal cell types. Dashed circles represent the cell bodies labeled by the respective marker. **(D, E)** Bar graphs representing the fraction of horizontal cells per type per retinal area. Data is given as mean ± standard deviation; n = 4 retinas from 3 animals, except for VT and DT in (D) where N = 3 retinas from 2 animals. **(F, G)** Distribution of nearest neighbor distances for each of the four horizontal cell types in the central (F) and peripheral chicken retina (G). The black and red vertical lines spanning the entire box give the median and the lower and upper quartiles of the nearest neighbor distances, respectively. NNRI represents the ratio between the mean of the nearest neighbor distance and the standard deviation of the cellular array (NNRI = nearest neighbor regularity index). Values above 1.9 are considered regular (Cook, 1996). Scale bars: A: 10 µm; C: 20 µm; inset: 10 µm.

For immunostainings of dye-injected slices and flat-mounts, specimens were washed thrice for 15 min in PBS and blocked in 5% donkey serum, 0.5% TritonX-100 and 0.02% sodium azide in PBS overnight at 4°C. Stainings were performed like stainings of vertical sections except that primary and secondary antibodies (Table 1) were applied in blocking solution for three days and overnight, respectively. After extensive washing, dye-injected slices were mounted in Mowiol (#81381, Sigma-Aldrich) with DAPI and flat-mount retinas in Vectashield (H-1000-10, Vector Labs).

### 2.4 Confocal image acquisition and analysis

Images were acquired with a confocal laser scanning microscope (Leica TCS SP8), using the HC PL APO 63×/1.4 or HC PL APO 40×/1.3 oil-immersion objectives. Zoom, pixel number, and z-step size were adjusted with respect to the experimental question. Stacks from dye-injected horizontal cells were deconvolved with Huygens Essential software (Scientific Volume Imaging, Hilversum, Netherlands; RRID:SCR_014237) using a theoretical point spread function. Images are presented as single optical sections or as maximum intensity projections of image stacks. Some confocal stacks were analyzed with Fiji (RRID:SCR_002285; Schindelin et al., 2012); the background was adjusted using the *Subtract Background* function and intensities were normalized using the *Contrast Enhancement* function. Images were occasionally filtered and/or pixels mildly saturated for presentation purposes.

To quantify horizontal cells in the flat-mounted chicken retina, six to eight images per flat-mount were taken, with at least one image in the periphery and central region of each quadrant (dorsal-nasal, DN; dorsal-temporal, DT; ventral-nasal, VN; ventral-temporal, VT). The *Cell Counter* plugin in Fiji was used to count the number of horizontal cells per type in an area of 291×291 µm and obtain their coordinates. A custom-written MATLAB 2023a (RRID:SCR_001622) script was used to perform a nearest neighbor analysis. First, we computed the distances between all cells of a given type and then filtered for the minimal distance for each cell. These nearest neighbor distances were pooled across all observed regions of the retina and sorted into 1 µm wide bins for the histogram. Further, the Nearest Neighbor Regularity Index (NNRI) for each cell type was computed by dividing the average nearest neighbor distance by their standard deviation (Cook, 1996).

### 2.5 Horizontal cell reconstruction in existing ssmSEM dataset

Our analysis of the horizontal cell type connectivity is based on the serial sectioning multi-beam electron microscopic (ssmSEM) dataset from Günther et al. (2024) (https://webknossos.mpinb.mpg.de/links/1GsNiZcF0C3fJjJC). The dataset originates from the dorsal periphery of a European robin retina and includes all layers of the retina in an area of 1 mm x 36.4 µm with a resolution of 4 nm x 4 nm x 40 nm. We identified 171 horizontal cell somata in the complete dataset. Since the dataset is restricted in z-depth due to limits in number of cut sections, we only reconstructed horizontal cells with their soma in the center of the z-plane to maximize the chances of reconstructing complete cells. Skeletonizations, volume reconstructions and subsequent synapse annotations were performed with the software webKnossos (Boergens et al., 2017). Unfortunately, all reconstructed cells were partial cells with different extents, but it is the largest ssmSEM dataset currently existing from any avian retina. We excluded cells with more than 10 dendrites or major parts of the dendritic field exceeding the volume of the dataset from further connectivity analysis.

Analysis of the synaptic connectivity of horizontal cells to photoreceptors was performed with modified Matlab2023b code from Günther et al. (2024). Since all cells were partial cells, we did not calculate dendritic tree diameters and absolute numbers of synapses per horizontal cell. Instead, we calculated the fraction of synapses to the individual photoreceptor cell types and the median number of synapses/photoreceptor/terminal. To further distinguish all single cone types, we measured the length of the primary axon, the diameter and position of the photoreceptor terminal in the outer plexiform layer, the position of the soma and the volume and position of the oil droplet. The length of the axon alone was already sufficient to separate blue-sensitive single cones from UV-sensitive single cones but for the group of red-/green-sensitive single cones, the terminal area in combination with oil droplet volume or position was necessary to discriminate the two types.

## 3 Results

### 3.1 Labeling different horizontal cell types in the chicken retina

To resolve the number of horizontal cell types in the chicken retina, we followed Fischer and colleagues (2007) and labeled vertical sections of the chicken retina for Calretinin (Calret), Islet1, and γ-aminobutyric acid (GABA). With this combination, we were able to distinguish four different types of horizontal cells in the inner nuclear layer (INL; Fig. 1A), which we termed H1 to H4 based on cell abundance and Mariani (1987). H1 cells were Calret+/GABA+ but Islet1- (blue arrowhead); H2 cells were Calret-/GABA-/Islet1+ (red arrowhead); H3 cells were Calret-/GABA+/Islet+ (brown arrowhead), and H4 cells were Calret+/GABA+/Islet1+ (yellow arrowhead). Using the same markers, we also labeled chicken retina flat-mounts and were able to distinguish the same types (Fig. 1B, C). This allowed us to study the distribution of horizontal cell types across the chicken retina by counting the number of cells per type in the peripheral and central chicken retina (Fig. 1B-G). H1 and H2 cells were similarly abundant (Fig. 1D-G; H1 cells: 48.6 ± 0.1% across all central quadrants; H2 cells:

39.8 ± 0.03% across all central quadrants; data always given as mean ± standard deviation of the mean); the same applies to H3 and H4 cells (Fig. 1D, E; H3 cells: 6.9 ± 0.02% across all central quadrants; H4 cells: 4.8 ± 0.01% across all central quadrants). Notably, the fraction of each type was rather constant across the visual field, i.e., in the retinal quadrants, and largely independent from retinal eccentricity (Fig. 1D, E). As expected, cell density was higher in the central than in the peripheral retina, ranging from 23,300 ± 1,900 horizontal cells/mm^2^ (n = 3 retinas from 2 chickens) in the central dorsal-temporal retina to 9,700 ± 1,300 horizontal cells/mm^2^ in the peripheral ventral-temporal retina (n = 4 retinas from 3 chickens).

A nearest neighbor analysis was performed to test the hypothesis of four different types based on the immunohistochemical marker profile. If cells belong to a distinct cell type, they are expected to “tile” the retina and show a defined nearest neighbor distance. Indeed, our analysis confirmed that four types of horizontal cells can be differentiated by Calret, GABA, and Islet-1 labeling in the chicken retina. All nearest neighbor distributions showed an “exclusion” zone in which no soma of the same type was found (Fig. 1F, G). In the central retina, H1 and H2 cells showed median nearest neighbor distances of 8.8 µm and 9.3 µm, respectively, whereas the nearest neighbor distances were much higher in H3 and H4 cells (20.4 µm and 27 µm, respectively; Fig. 1F). A similar difference was seen for the peripheral retina (median nearest neighbor distance: H1: 11 µm; H2: 10.7 µm; H3: 27.7 µm; H4: 40.3 µm; Fig. 1G). This is in line with the lower density of H3 and H4 cells compared to H1 and H2 cells (Fig. 1D, E). In addition, we also calculated the nearest neighbor regularity index (NNRI) which represents the average nearest neighbor distance divided by its standard deviation. True retinal mosaics consistently yield NNRI values >2.0 whereas an NNRI of 1.91 shows complete randomness (Cook, 1996; Keeley et al., 2017). All horizontal cell types had NNRI >2.7 (Fig. 1F, G).

### 3.2 The connectivity of chicken horizontal cells

To study the connectivity of the different horizontal cell types of the chicken retina, we dye-injected individual horizontal cells in vibratome slices and subsequently labeled for PSD95, a marker for all photoreceptor terminals in the bird retina. Photoreceptor types were identified based on their stratification in the outer plexiform layer (OPL): double cones and rods terminate in the most distal layer, green and red cones in the middle and violet and blue cones in the most proximal layer of the OPL (Balaji et al., 2023; Günther et al., 2021; Mariani, 1987). To ease photoreceptor identification, horizontal cells were injected in the peripheral retina as photoreceptor terminals become larger and less numerous in this region (Balaji et al., 2023). H1 cells were identified based on their small and compact dendritic field and the axon protruding from the soma; they seemed to contact all cone photoreceptors within reach (Fig. 2). Whether or not they also contact the rather small rod terminals in the most distal part of the OPL, we could not resolve with this approach. H2 cells were identified by their much larger dendritic field and their sparse branching. These cells almost exclusively contacted double cone photoreceptors, with a clear preference for the accessory member (Fig. 3A, asterisks), as reported earlier (Tanabe et al., 2006). The process marked with an arrow represents a dendritic branch which makes a sharp turn and leaves the confocal stack. As H3 and H4 cells are very sparsely distributed in the retina, especially in the periphery, we were only able to inject a single cell that showed a morphology distinct from H1 and H2 cells. The cell showed a very flat and large dendritic arbor and contacted putative green and/or red cones which stratify in the middle of the OPL (Fig. 4). However, there are also processes reaching out to violet/blue cones (Fig. 4B, circle). As these processes do not invaginate the terminals, we could not determine whether these are true contacts and whether this cell belonged to type H3 or H4 cells. However, the large dendritic tree confirmed earlier reports (Mariani, 1987; Tanabe et al., 2006) and is in line with the lower density and larger nearest neighbor distances (Fig. 1D-G) we found for H3 and H4 cells.

**Figure 2.**
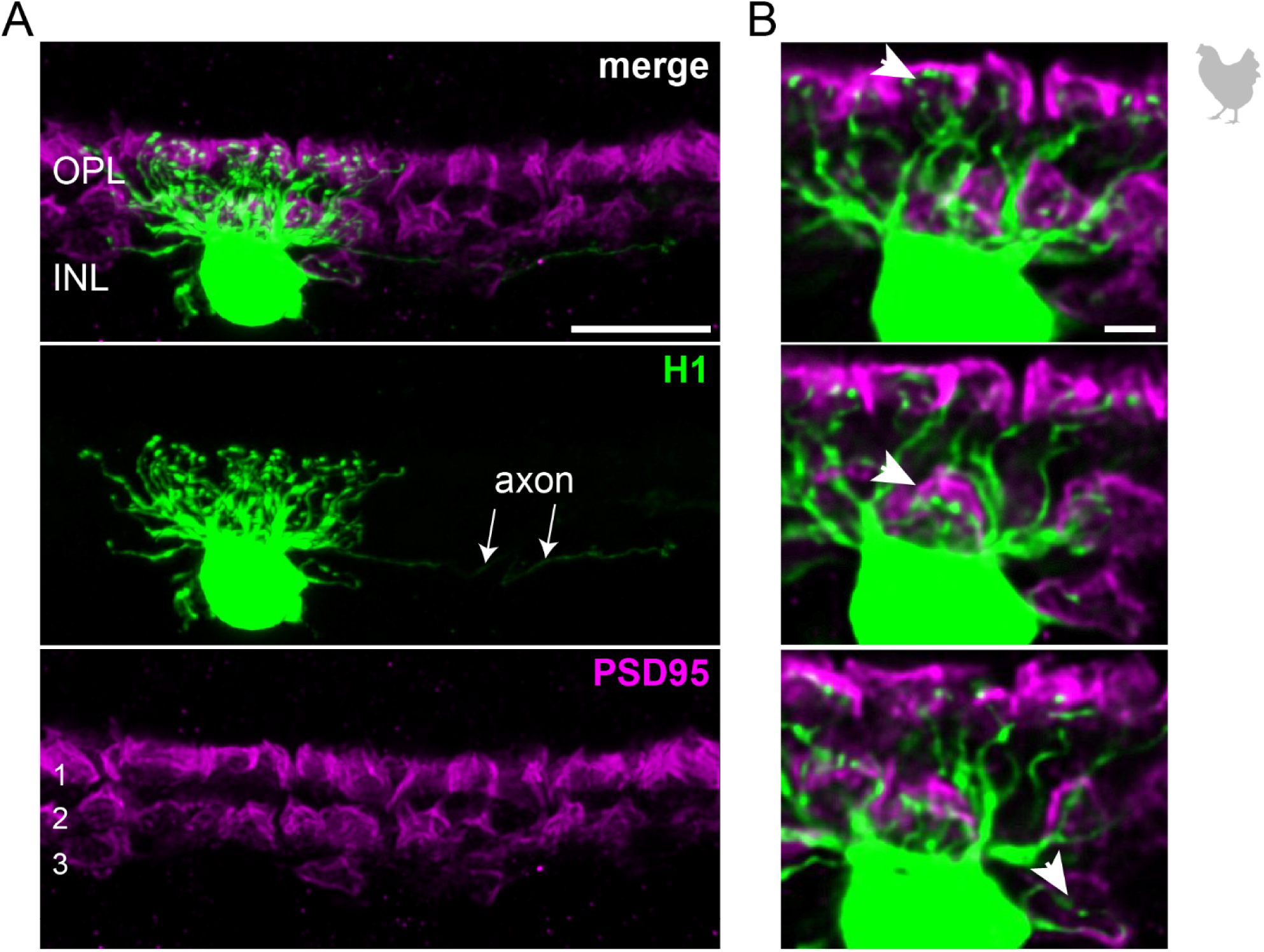
The axon-bearing H1 horizontal cell in the peripheral chicken retina contacts all cones. **(A)** Maximum projection of a dye-injected H1 horizontal cell, revealing a dense and narrow dendritic field and an axon (arrows). Double labeling with PSD95 reveals the stratification of photoreceptor terminals in three layers (labeled 1-3) of the outer plexiform layer (OPL). Please note that for PSD95, the maximum projection of a substack is shown to better illustrate the three layers. **(B)** Maximum projections of substacks (5-10 optical sections) at different positions of the confocal stack. H1 cell dendrites contact cones in all three layers of the OPL (arrowheads). Whether or not rods are also contacted by the dendritic field could not be resolved. Scale: 10 µm (A), 2 µm (B).

**Figure 3.**
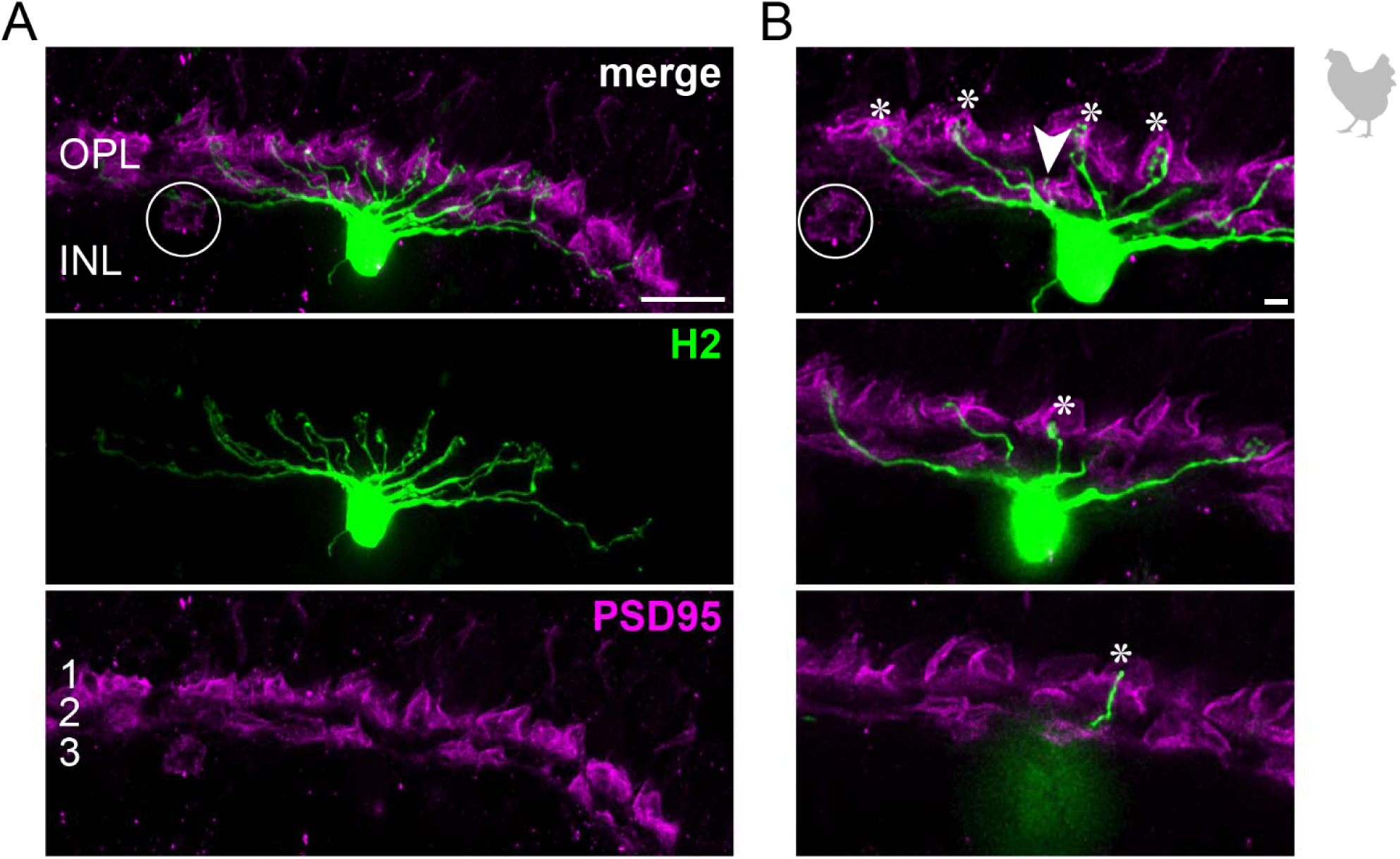
H2 horizontal cells in the peripheral chicken retina make selective contacts to the accessory member of the double cone photoreceptors. **(A)** Dye-injected H2 horizontal cell, revealing a large dendritic tree which is only branching in the terminal endings. A maximum projection is shown. Double labeling with PSD95 reveals the stratification of photoreceptor terminals in three layers (labeled 1-3) of the outer plexiform layer (OPL). Please note that for PSD95, the maximum projection of a substack is shown to better illustrate the three layers. **(B)** Maximum projections of substacks (12-18 optical sections). The vast majority of H2 cell dendrites contact the accessory members of the double cones in the most distal OPL layer (1, asterisks), while avoiding photoreceptor terminals in layer 3 belonging to violet and blue cones, marked by a circle. However, one dendrite seems to contact a green or red cone in layer 2 of the OPL (arrowhead). Scale: 10 µm (A), 2 µm (B).

**Figure 4.**
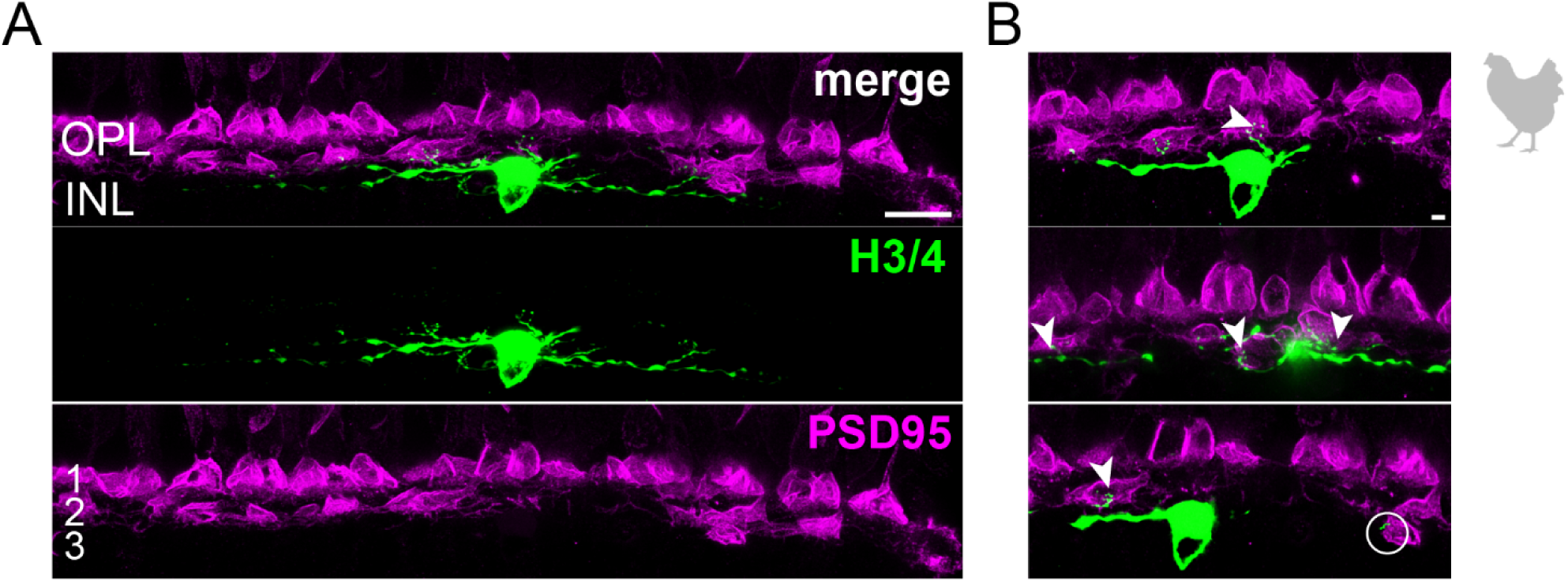
Wide-field horizontal cell (H3 or H4) in the peripheral chicken retina avoiding the outermost layer of the outer plexiform layer. **(A)** Maximum projection of a dye-injected wide-field horizontal cell, revealing a flat dendritic field with processes mostly confined to layer 2 of the outer plexiform layer. Whether this cell represents an H3 or H4 cell is not possible to discern. Double labeling with PSD95 reveals the stratification of photoreceptor terminals in three layers (labeled 1-3) of the outer plexiform layer (OPL). Please note that for PSD95, the maximum projection of a substack is shown to better illustrate the three layers. **(B)** Maximum projections of substacks (30-35 optical sections) of the confocal stack. H3/4 cell dendrites contact mostly red and/or green cones in layer 2 of the OPL (arrowheads), while avoiding rods and double cones. Rarely, they reach out to violet/blue cones (circle). However, whether these dendrites which do not invaginate the cone pedicle represent true contacts is unclear. Scale: 10 µm (A), 2 µm (B).

In summary, our immunostainings confirm four different types of horizontal cells in the chicken retina which can be distinguished based on their immunoreactivity profile and show distinct connections to photoreceptors.

### 3.3 Labeling different horizontal cell types in the European robin retina

Next, we used the same markers (Calret, Islet1, GABA) to analyze horizontal cell types in the retina of the European robin, a night-migratory and flight-hunting songbird with a light-dependent magnetic sense (Wiltschko et al., 1993; Xu et al., 2021; Zapka et al., 2009) whose foraging behavior is very different from ground-pecking birds, such as the chicken (Fischer et al., 2007) and the pigeon (Mariani, 1987). In retinal cryosections of the European robin retina, we resolved three different types of horizontal cells based on this approach (Fig. 5): H1 cells were Calret+/GABA+ but Islet1- (blue arrowhead); H2 cells were Calret-/GABA-/Islet1+ (red arrowhead); H3 cells were Calret-/GABA+/Islet+ (dark-red arrowhead). We did not find cells that were positive for all markers (potential H4 cells), presumably because of the weak GABA labeling in retinas of the European robin.

**Figure 5.**
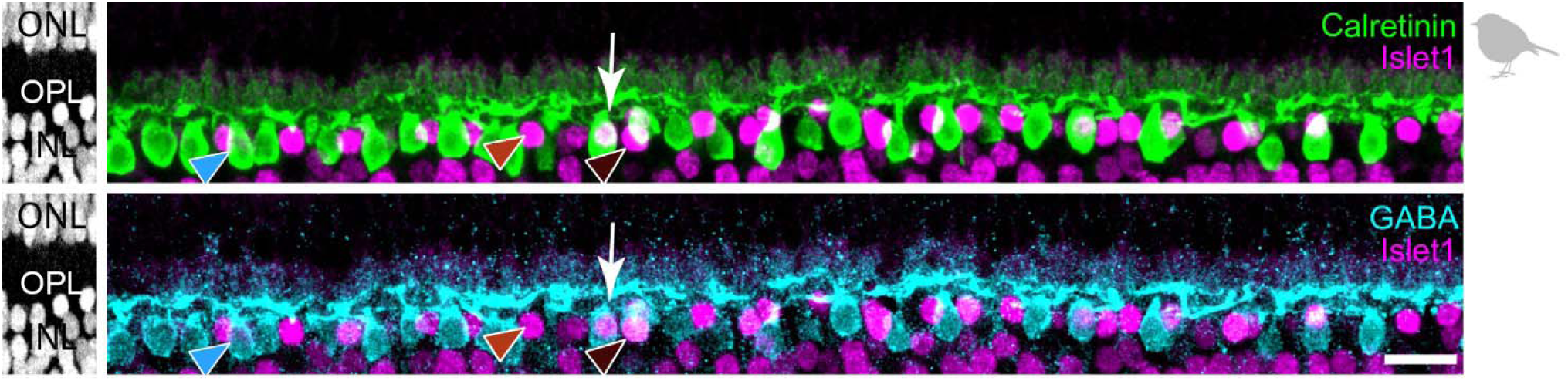
Triple labeling with Calretinin, GABA, and Islet1 allows to differentiate only three types of horizontal cells in the European robin retina. Maximum projections of vertical slices of the European robin retina labeled for Calretinin (Calret), GABA, and Islet1. Based on these markers, three different types of horizontal cells were distinguished: H1: Calret+/GABA+ (blue arrowhead); H2: Islet1+ (red arrowhead); H3: Islet+/GABA+ (brown arrowhead). The white arrow points to a cell which seems to express all three markers, but inspection of the confocal stack showed that an Islet1+ H2 cell lies behind a Calret+/GABA+ H1 cell. Scale bar: 10 µm.

### 3.4 The connectivity of robin horizontal cells

To further investigate the number of horizontal cell types and their synaptic connectivity to photoreceptors in the European robin retina, we used the publicly available serial-sectioning multi-beam EM dataset from the dorsal periphery of the European robin retina (Günther et al., 2024) and reconstructed dendritic and axonal arbors of horizontal cells. In total, we identified 171 horizontal cell somata in the INL, from which we reconstructed 65 horizontal cells. Based on their morphology, we grouped the cells into four types: H1 (N=35), H2 (N= 21), H3 (N=6), and H4 (N=3) horizontal cells, again following the nomenclature of Mariani (1987) (Fig. 6A). Of the 65 reconstructed horizontal cells, 31 (14 H1, 9 H2, 5 H3, and 3 H4 cells) were further synapse annotated. This analysis revealed that the dendrites of each horizontal cell type connected selectively to a different set of cone photoreceptors (Fig. 6B, C, see details below) while avoiding rod terminals, as described for the pigeon retina (Mariani, 1987).

**Figure 6.**
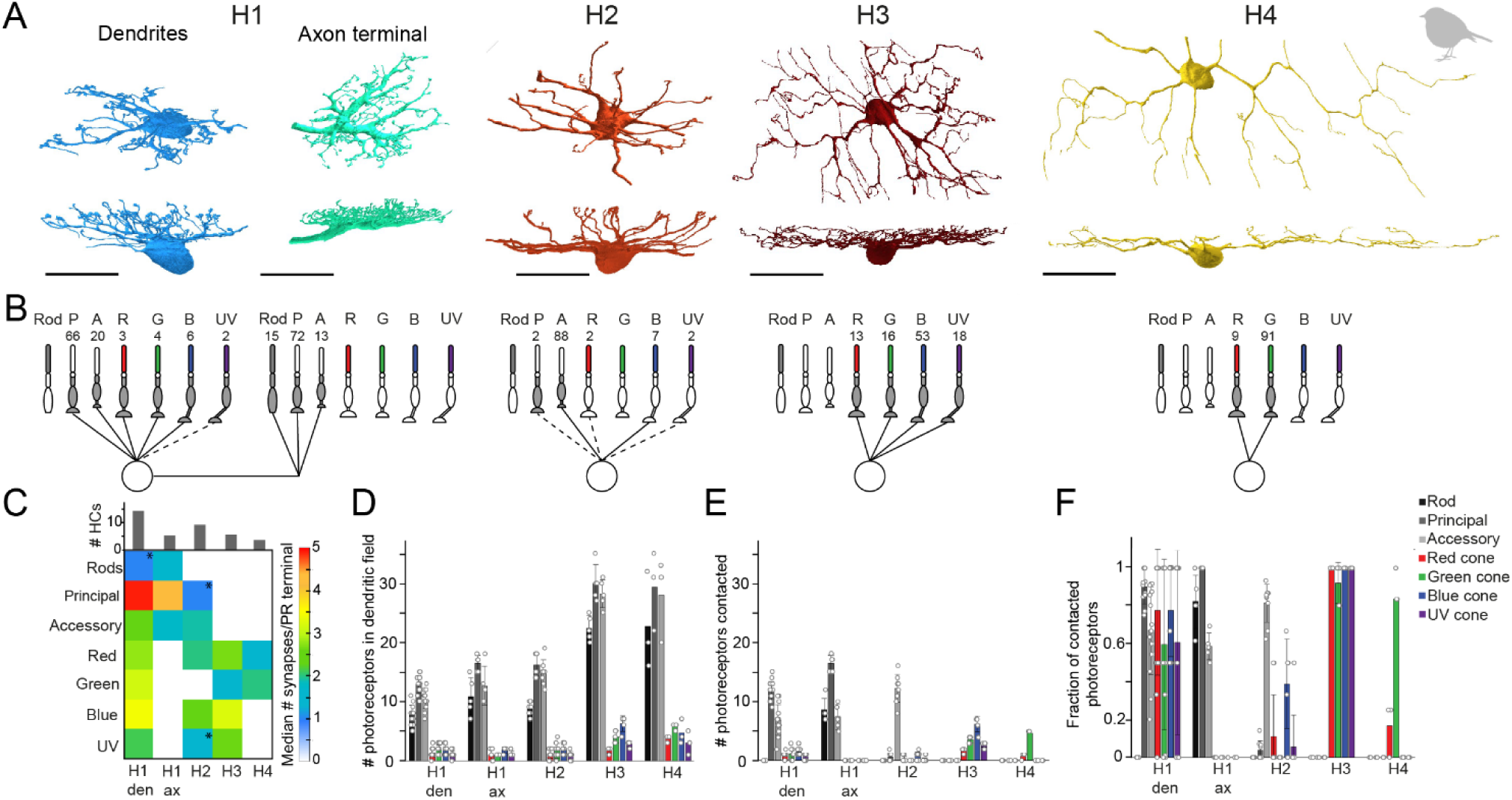
Quantification of horizontal cell type specific contacts to photoreceptors. **(A)** Volume reconstructions of each horizontal cell type. In total, we identified four horizontal cell types namely H1 (dendrites N = 35, axons N = 5), H2 (N = 21), H3 (N = 6), and H4 (N = 3). All scale bars: 20 µm. **(B)** Network motifs for each horizontal cell type. The numbers above each photoreceptor type represent the fraction of synapses/photoreceptor type in percent. Dashed lines indicate that this photoreceptor cell type was rarely (<3%) contacted by the respective horizontal cell. P = principal member of a double cone, A = accessory member of a double cone, R = red single cone, G = green single cone, B = blue single cone, UV = ultraviolet single cone. **(C)** Connectivity matrix with the median number of synapses/photoreceptor (PR) terminal per horizontal cell type. Histograms above the connectivity matrix indicate the number of cells that were included in the analysis. Connections between horizontal cells and photoreceptor cells that are highlighted by an asterisk only occurred in a few individual horizontal cells; den = dendrites; ax = axon terminal. **(D)** Mean number ± standard deviation (SD) of rod and cone terminals within the dendritic field of different horizontal cells. **(E)** Mean number ± SD of contacted photoreceptor cells/type for different horizontal cell types. **(F)** Mean fraction ± SD of contacted photoreceptors from photoreceptors in the dendritic field of different horizontal cell types.

To investigate whether rod photoreceptors are contacted by the axon terminals of horizontal cells, as reported for axon-bearing horizontal cells in the mammalian retina (Peichl and Gonzalez-Soriano, 1994; Trümpler et al., 2008), we reconstructed cells starting at the lateral elements of the rod ribbon synapses. Only the trees with a prominent axon-like fiber were considered axonal trees (N = 5; Fig. 6A). Since H1 cells are the only horizontal cells described to bear an axon (Mariani, 1987) and were the only horizontal cells we found during reconstructions to bear a prominent axon-like fiber (exceeding the limits of the dataset), we presume that the reconstructed axon terminals belong to H1 cells.

H1 horizontal cells received frequent input from all types of cones within their dendritic field (Fig. 6D-F) but formed most synapses with the principal member of double cones (Fig. 6B). This results from the high number of contacts per terminal (Fig. 6C) and the higher abundance of double cones in comparison to single cones (Fig. 6D). H1 axon terminals contacted rod terminals but also double cone photoreceptors with a strong preference for the principal member of the double cone (Fig. 6B), consistent with a previous report on chicken (Tanabe et al., 2006). In contrast, H2 horizontal cells were very selective for the accessory member of the double cone (93.6% of all contacts) (Fig. 6B) but with a comparatively low number of synapses per terminal (Fig. 6C). H3 and H4 horizontal cells are both single cone selective horizontal cells but differ greatly in the type of single cones they contact (Fig. 6B). H3 horizontal cells contacted all single cones within their dendritic field, irrespective of the type (Fig. 6D-F). However, due to the higher abundance of blue cones combined with a higher number of contacts per terminal to blue cones, H3 horizontal cells formed more than half of their synapses with blue cones. In contrast, H4 horizontal cells almost exclusively contacted green cones (Fig. 6B-F). Please note that the preference of each horizontal cell type for a specific photoreceptor type is the combination of contacted photoreceptors terminals (Fig. 6D-F) and the number of contacts within one terminal (Fig. 6C). Most horizontal cells with a preference for a specific photoreceptor type not only contacted these terminals more frequently (Fig. 6F) but also made more contacts within the respective terminal. H2 cells seem to be an exception to this rule since they generally only made a low number of contacts to any contacted terminal. As a result, the preference of H2 cells to accessory members purely results from the absolute number of contacted terminals.

During horizontal cell reconstructions, we identified conventional chemical synapses at the dendrites and somata of specific horizontal cells (Fig. 7A, B). A quantitative analysis revealed that these synapses are almost exclusively located on somata and dendrites of H1 horizontal cells; we only found two very small chemical synapses on the dendrites of H3 horizontal cells (Fig. 7C, D). Following up on this finding, we also screened the chicken EM dataset (Günther et al., 2021) for conventional synapses at the somata or dendrites of horizontal cells and found similar synapses (N=18 on 3 horizontal cells) that were also exclusively located on H1 horizontal cells (data not shown). To investigate the specificity of these H1 cell synapses in the European robin retina, we reconstructed the postsynaptic partners. Most contacts were formed with type B7 bipolar cells (Günther et al., 2024), suggesting a feedforward pathway selective for a certain bipolar cell type. In addition, we found conventional synapses with previously undescribed cell types (Fig. 7) and (partially) reconstructed the cell whose dendritic and axonal field was the most complete (Fig. 7E). The axonal field did not contain ribbon synapses, which would be characteristic of a bipolar cell, but conventional synapses, indicative of an amacrine cell. Additionally, the dendrites of this cell barely contacted photoreceptor terminals but formed conventional synapses in the OPL. Reconstructing the postsynaptic partners of those conventional synapses revealed that most of the postsynaptic partners were B4a bipolar cells. This circuit analysis suggests that the avian retina contains an interplexiform cell which receives chemical input from H1 horizontal cell dendrites and provides output to B4a bipolar cells.

**Figure 7.**
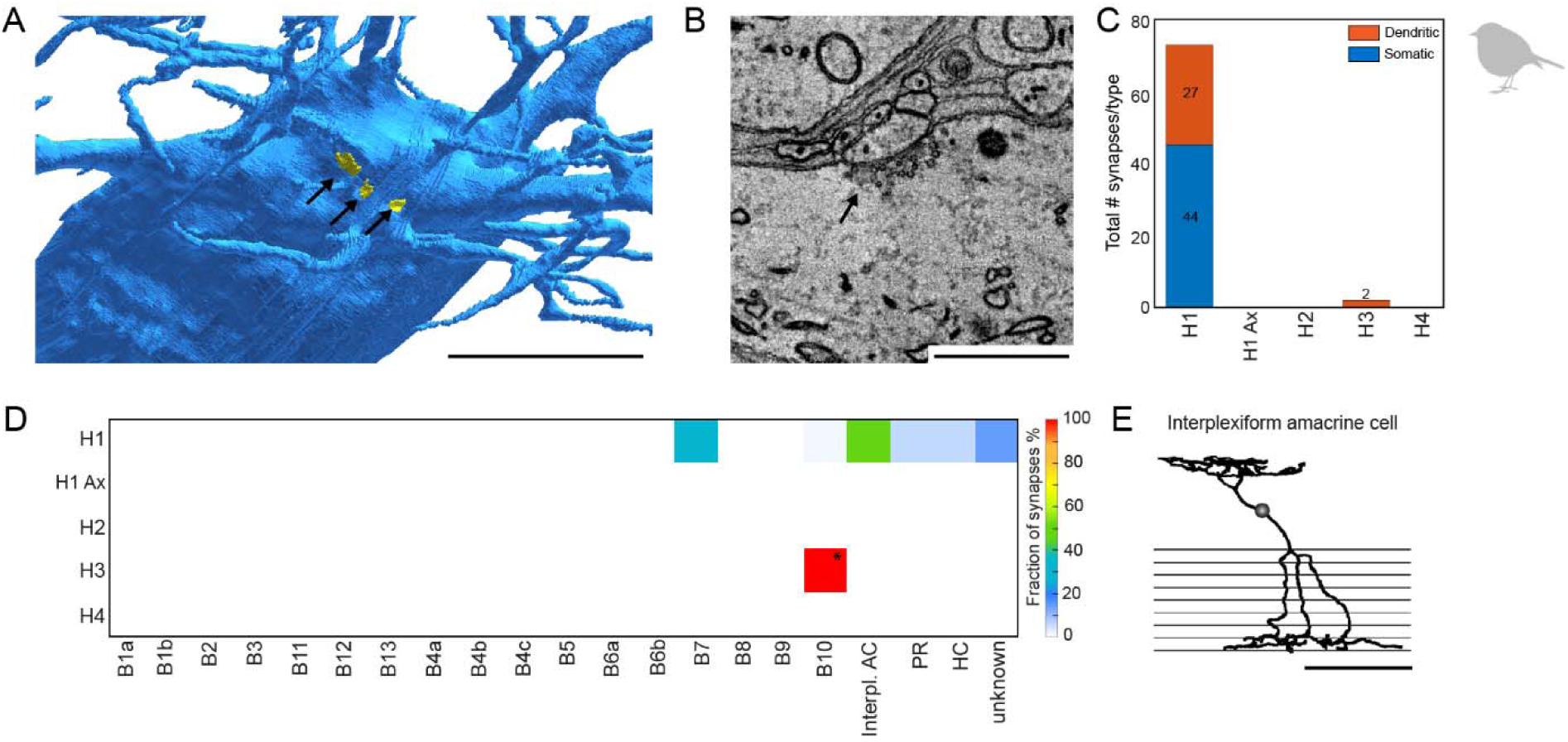
H1 horizontal cells form conventional synapses at their dendrites and somata to specific bipolar and interplexiform amacrine cells. **(A)** Volume reconstructed soma of an H1 horizontal cell (blue) with arrows pointing to the locations of conventional synapses highlighted in yellow. Scale bar: 5 µm. **(B)** Electron microscopic image of a conventional synapse at the soma of a H1 cell. Scale bar: 1 µm. **(C)** Total number of conventional synapses for each HC type separated by their location either on the cell body or the dendrites. In total, we inspected 21 H1, 18 H2, 6 H3 and 3 H4 horizontal cells. **(D)** Connectivity matrix with fractionated number of synapses for each postsynaptic cell type in percent. Only two small synapses were found at H3 horizontal cell dendrites (marked with *). Postsynaptic cells from the H3 horizontal cell synapses resembled the previously described B10 bipolar cell in the chicken (Günther et al. 2021). Ax = axon terminal; B1-B10, bipolar cell types; interpl. AC, interplexiform amacrine cell; HC, horizontal cell; PR, photoreceptor. **(E)** Skeletonized postsynaptic interplexiform amacrine cell, stratifying in the outer and inner plexiform layer. Scale bar: 50 µm.

## 4 Discussion

We first used an immunohistochemical approach to 1) confirm horizontal cell types in the chicken retina and 2) expand the analysis on previously undescribed European robin horizontal cells. In a second approach, we used the previously published electron microscopic dataset from the peripheral European robin retina to 3) identify the detailed connectivity between photoreceptor cell types and each individual horizontal cell type and 4) provide first insights into potential locations of synapses for feedforward inhibition.

### 4.1 Bird retinas contain four types of horizontal cells

Horizontal cells had so far only been studied in two bird species, the chicken (Edqvist et al., 2008; Fischer et al., 2007; Tanabe et al., 2006) and the pigeon (Mariani, 1987) where either three (Edqvist et al., 2008; Tanabe et al., 2006) or four types (Fischer et al., 2007; Mariani, 1987) were differentiated, depending on the choice of method. Here, we confirmed four types of horizontal cells in the chicken retina using similar markers as Fischer et al. (2007). However, while their types H3 and H4 were GABA- (Fischer et al., 2007), we show that these two types are GABA+. We also studied the distribution of the four types across the chicken retina by probing four central and four peripheral regions in each retinal quadrant. Each horizontal cell type made up a similar proportion of all horizontal cells across the retina, with differences only in the central dorsotemporal retina, where H2 cells were less abundant and H3 cells slightly more abundant than in other quadrants. Across all quadrants, H1 and H2 cells were the major horizontal cell types whereas H3 and H4 cells were both rather rare as was reported before (Fischer et al., 2007) (for numbers, see results section). Whether the horizontal cell density in the chicken retina is proportional to the density of photoreceptors and shows regional hot spots as reported for the mouse retina (Spinelli et al., 2024) remains elusive as this would require much denser sampling of the entire bird retina. Given the occasionally weak penetration of the antibodies, we refrained from such an analysis. In our nearest neighbor analysis, H1-4 cells showed marked exclusion zones and an NNRI > 2.7 for all horizontal cell types, suggesting that each type forms its own retinal mosaics (Cook, 1996; Keeley et al., 2017). This is also consistent with recent transcriptomic data, which indicates four horizontal cell types in the chicken (Yamagata et al., 2021).

The same antibody combination (Calret, GABA, and Islet1), however, yielded only three different types of horizontal cells in the European robin retina, which was probably caused by the low penetration of the GABA antibody. To overcome this obstacle, we made use of a serial electron microscopy stack and were able to confirm four types also for this species. The reconstructed horizontal cells in the European robin retina are morphologically very similar to the different types identified in the chicken (this study; Fischer et al., 2007) and pigeon retina (Mariani, 1987). This is consistent with the notion that the outer retina is evolutionary more conserved than the inner retina (Baden, 2024; Hahn et al., 2023), which in case of birds already shows marked differences at the level of bipolar cells (Balaji et al., 2023; Günther et al., 2024, 2021).

### 4.2 Bird horizontal cells make type-specific contacts with photoreceptors

We also used the high-resolution information of the serial EM stack to fully resolve the horizontal cell/ photoreceptor connectivity in the dorsal peripheral retina of the European robin. Intracellular dye injections into chicken horizontal cells complemented this approach. Both approaches show that H1 cell dendrites predominantly contact the double cones, with a strong preference for the principal member. H1 dendrites also contact single cones, while avoiding rod terminals. The axon terminals of H1 cells preferentially synapse with the principal member of the double cone but also contact the accessory member and rods. A similar connectivity was suggested for H1 cells in the pigeon retina (Mariani, 1987). For H2 cells, our data confirms the preferential connection to the accessory member of the double cone (Tanabe et al., 2006), again demonstrating that both members of the double cone make independent connections with horizontal and bipolar cells (Baden, 2024; Günther et al., 2024, 2021). Although there were some doubts on the existence of a fourth horizontal cell type (Edqvist et al., 2008; but see Fischer et al., 2007), analyzing the photoreceptor connectivity yielded a clear result: H3 cells contact all single cones within their reach (with a preference for blue cones which were more abundant in the dorsal peripheral retina), whereas H4 preferentially contact green cones.

### 4.3 Similarities with the turtle retina allow predicting the putative light responses of bird horizontal cells

Horizontal cells are crucial for making color opponent computations in the retina (reviewed in Thoreson and Dacey, 2019). The horizontal cell types and their photoreceptor connections found here in birds are strikingly similar to those described in the turtle retina, which also contains a similar set of photoreceptor types (Bowmaker, 2008). As horizontal cells in the turtle retina were intensely studied, we speculate here on the function of bird horizontal cells. Like the bird retina, the turtle retina contains four different types of horizontal cells, one axon-bearing B-type and three axon-less A-type horizontal cells (Gallego, 1986; Leeper, 1978). The axon-bearing turtle H1 cell (corresponding to the H1 cell described here) was shown to contact double and single cones (Leeper, 1978). It is a luminosity-type horizontal cell as it lacks color opponency and hyperpolarizes in response to all wavelengths of light (Tranchina et al., 1983). Thus, it seems likely that bird H1 cells, which also sample from double and single cones like turtle H1 cells, also belong to the luminosity type and encode contrast rather than spectral information. Bird H2 cells may correspond to turtle H4 cells. Both horizontal cell types preferentially connect to the double cone accessory member (Leeper, 1978) which expresses red opsin in both, birds and turtles (Davies et al., 2012; Loew and Govardovskii, 2001). Both cells may correspond to a second luminosity type although the physiological responses of turtle H4 cells are not entirely clear (Ammermüller and Kolb, 1996), presumably due to their low abundance. Bird H3 cells may be counterparts of turtle H3 cells as both cell types show a preference for short-wavelength input (Kolb et al., 1988). Turtle H3 cells respond biphasically to light stimuli, depolarizing to red light and hyperpolarizing to blue (Ammermüller et al., 1995). They therefore belong to chromaticity horizontal cells, which may also be the case for bird H3 cells. Bird H4 cells prefer green cones and may consequently correspond to turtle H2 cells, which wire to blue and green cones (Leeper, 1978) and show color-opponent responses, i.e. depolarization to red and hyperpolarization to green (Ammermüller et al., 1995). In summary, in the bird retina, the two most abundant horizontal cell types (H1 and H2) may encode intensity whereas the two least abundant cell types (H3 and H4 cells) may show color opponency. Recordings from bird horizontal cells, which have not been described yet, will show whether these predictions are correct.

### 4.4 Potential horizontal cell involvement in processing light-dependent magnetic information

The double cones of European robins contain the blue-light absorbing and magnetically sensitive protein, Cryptochrome 4 (Chetverikova et al., 2022; Günther et al., 2018; Xu et al., 2021), which is thought to be the primary magnetic sensor in light-dependent, radical-pair-based magnetoreception (Hore and Mouritsen, 2016; Mouritsen, 2018). Furthermore, the accessory member of the double cone in the European robin lacks the oil droplet (Günther et al., 2024) that filters out blue light in some other bird species (Bowmaker et al., 1997). Therefore, the H2 cell of the European robin, which contacts almost exclusively the accessory member of the double cone, may carry magnetic information.

Horizontal cell-mediated inhibition may create antagonistic receptive fields at the level of bipolar cells which are then passed on to ganglion cells (Ströh et al., 2018). Therefore, it is also conceivable that H1 cells, which contact almost all photoreceptors, play a role in magnetoreception. They could modulate type 6 bipolar cells (Günther et al., 2024) which sample exclusively from accessory members of the double cone. The potential combination of selective sampling (type 6 bipolar cells) and non-selective sampling (H1 cells) could generate a center-surround pathway for magnetic signals, comparable to the color-opponent pathways in the peripheral primate retina (Field et al., 2010).

### 4.5 Potential sites for feedforward inhibition in bird horizontal cells

Horizontal cells shape the receptive fields of downstream neurons by feedback and feedforward signaling to photoreceptors and bipolar cells, respectively. Only recently, the synaptic sites for feedforward inhibition were identified in the mouse retina (Behrens et al., 2021) where they occurred as short segments of increased dendritic diameter on the primary dendrites of horizontal cells. Here, we also identified conventional synapses in horizontal cells, which predominantly occurred on the somata of H1 cells. These structures showed synaptic vesicles (Fig. 7B) which may contain GABA as H1 cells are GABA+ (Fischer et al., 2007; this study). Contacts were made with 1) type 7 bipolar cells suggesting that feedforward inhibition is bipolar cell type specific (Behrens et al., 2021); 2) other horizontal cells (as was suggested before Dowling et al., 1966); and 3) an interplexiform cell which presumably provides output to type 4a bipolar cells and stratifies in layer 8 of the INL (Fig. 7E). The latter cell did not resemble any of the interplexiform cells reported earlier for the bird retina (Kalloniatis and Fletcher, 1993; Mariani, 1987) and the physiological function of these connections is unclear. In conclusion, our data provides a first insight into the synaptic complexity of the bird OPL where horizontal cells not only form feedback synapses with photoreceptors but also feedforward synapses with specific types of bipolar cells and at least one type of interplexiform cells.

In summary, two distantly related bird species each contain four types of horizontal cells, suggesting that this is a conserved feature of avian species. Bird horizontal cells show similar connections to photoreceptors as the ones in reptiles. Based on this, we speculate that bird retinas may contain both intensity-encoding and color-opponent horizontal cells, corresponding to H1/H2 and H3/4, respectively. Electrophysiological studies are needed to test for the functional role of horizontal cells in the superb spatial and color vision of birds.

## 5 Conflict of Interest

The authors declare that the research was conducted without any commercial or financial relationships that could be construed as a potential conflict of interest.

## 6 Author Contributions

KD, HM, and AG conceived the study and designed experiments; AG, VB, and JJF performed experiments; AG, VB, JJF, BL, AYR, TW and DA analyzed data; all contributed to the interpretation of the data; AG, VB, KD, and JJF created figures; AG and KD wrote the manuscript; all authors edited and commented on the manuscript and KD finalized it.

## 7 Funding

This project was funded by the Deutsche Forschungsgemeinschaft (DFG; Walter Benjamin Fellowship, project 508628753 to AG; SFB 1372: Magnetoreception and Navigation in Vertebrates, project 395940726 to HM and KD). HM also acknowledges funding from the European Research Council (under the European Union’s Horizon 2020 research and innovation programme, grant agreement no. 810002; Synergy Grant: QuantumBirds). In addition, funding was provided by the Max-Planck-Institute for Neurobiology of Behavior – caesar (to AG).

## 8 Acknowledgments

We thank Matteo Spinelli and Dr. Christian Puller for sharing their brushing technique with us. We also thank Leonie L. Pfeiffer and Irina Fomins for their excellent technical assistance and the administration team of the Animal Navigation group for their constant administrative support. We also thank Eric Jelli for his support with volume reconstructing the horizontal cells in the EM dataset. The authors also gratefully acknowledge the Core Facility Fluorescence Microscopy of the Carl von Ossietzky Universität Oldenburg, Oldenburg, Germany.

## 10 Data Availability Statement

The EM dataset used in this study was already published in (Günther et al., 2024) and can be found in the webknossos depository under: https://webknossos.mpinb.mpg.de/links/1GsNiZcF0C3fJjJC. The raw data supporting the conclusions of this article will be made available by the authors, without undue reservation.

## Notes

### Competing Interest Statement

The authors have declared no competing interest.

## References

Ammermüller, J., Kolb, H., 1996. Functional architecture of the turtle retina. Progress in Retinal and Eye Research 15, 393–433. 10.1016/1350-9462(96)00009-2

Ammermüller, J., Muller, J.F., Kolb, H., 1995. The organization of the turtle inner retina. II. Analysis of color-coded and directionally selective cells. J Comp Neurol 358, 35–62. 10.1002/cne.903580103

Asi, H., Perlman, I., 1998. Neural interactions between cone photoreceptors and horizontal cells in the turtle (Mauremys caspicd) retina. Visual Neuroscience 15, 1–13. 10.1017/s0952523898146047

Baden, T., 2024. The vertebrate retina: a window into the evolution of computation in the brain. Current Opinion in Behavioral Sciences 57, 101391. 10.1016/j.cobeha.2024.101391

Baden, T., Berens, P., Franke, K., Román Rosón, M., Bethge, M., Euler, T., 2016. The functional diversity of retinal ganglion cells in the mouse. Nature 529, 345–350. 10.1038/nature16468

Balaji, V., Haverkamp, S., Seth, P.K., Günther, A., Mendoza, E., Schmidt, J., Herrmann, M., Pfeiffer, L.L., Němec, P., Scharff, C., Mouritsen, H., Dedek, K., 2023. Immunohistochemical characterization of bipolar cells in four distantly related avian species. J Comp Neurol 531, 561–581. 10.1002/cne.25443

Behrens, C., Schubert, T., Haverkamp, S., Euler, T., Berens, P., 2016. Connectivity map of bipolar cells and photoreceptors in the mouse retina. Elife 5, e20041. 10.7554/eLife.20041

Behrens, C., Yadav, S.C., Korympidou, M.M., Zhang, Y., Haverkamp, S., Irsen, S., Schaedler, A., Lu, X., Liu, Z., Lause, J., St-Pierre, F., Franke, K., Vlasits, A., Dedek, K., Smith, R.G., Euler, T., Berens, P., Schubert, T., 2021. Retinal horizontal cells use different synaptic sites for global feedforward and local feedback signaling. Curr Biol S0960–9822(21)01641–9. 10.1016/j.cub.2021.11.055

Boergens, K.M., Berning, M., Bocklisch, T., Bräunlein, D., Drawitsch, F., Frohnhofen, J., Herold, T., Otto, P., Rzepka, N., Werkmeister, T., Werner, D., Wiese, G., Wissler, H., Helmstaedter, M., 2017. webKnossos: efficient online 3D data annotation for connectomics. Nat Methods 14, 691–694. 10.1038/nmeth.4331

Boije, H., Shirazi Fard, S., Edqvist, P.-H., Hallböök, F., 2016. Horizontal Cells, the Odd Ones Out in the Retina, Give Insights into Development and Disease. Front. Neuroanat. 10. 10.3389/fnana.2016.00077

Bowmaker, J.K., 2008. Evolution of vertebrate visual pigments. Vision Research, Vision Research Reviews 48, 2022–2041. 10.1016/j.visres.2008.03.025

Bowmaker, J.K., Heath, L.A., Wilkie, S.E., Hunt, D.M., 1997. Visual pigments and oil droplets from six classes of photoreceptor in the retinas of birds. Vision Research 37, 2183–2194. 10.1016/S0042-6989(97)00026-6

Chaya, T., Matsumoto, A., Sugita, Y., Watanabe, S., Kuwahara, R., Tachibana, M., Furukawa, T., 2017. Versatile functional roles of horizontal cells in the retinal circuit. Sci Rep 7, 5540. 10.1038/s41598-017-05543-2

Chetverikova, R., Dautaj, G., Schwigon, L., Dedek, K., Mouritsen, H., 2022. Double cones in the avian retina form an oriented mosaic which might facilitate magnetoreception and/or polarized light sensing. J R Soc Interface 19, 20210877. 10.1098/rsif.2021.0877

Connaughton, V.P., Nelson, R., 2010. Spectral Responses in Zebrafish Horizontal Cells Include a Tetraphasic Response and a Novel UV-Dominated Triphasic Response. Journal of Neurophysiology 104, 2407–2422. 10.1152/jn.00644.2009

Cook, J.E., 1996. Spatial properties of retinal mosaics: an empirical evaluation of some existing measures. Vis Neurosci 13, 15–30. 10.1017/s0952523800007094

Davies, W.I.L., Collin, S.P., Hunt, D.M., 2012. Molecular ecology and adaptation of visual photopigments in craniates. Mol Ecol 21, 3121–3158. 10.1111/j.1365-294X.2012.05617.x

Dowling, J.E., Brown, J.E., Major, D., 1966. Synapses of Horizontal Cells in Rabbit and Cat Retinas. Science 153, 1639–1641. 10.1126/science.153.3744.1639

Drinnenberg, A., Franke, F., Morikawa, R.K., Jüttner, J., Hillier, D., Hantz, P., Hierlemann, A., Azeredo da Silveira, R., Roska, B., 2018. How Diverse Retinal Functions Arise from Feedback at the First Visual Synapse. Neuron 99, 117–134.e11. 10.1016/j.neuron.2018.06.001

Edqvist, P.-H.D., Lek, M., Boije, H., Lindbäck, S.M., Hallböök, F., 2008. Axon-bearing and axonless horizontal cell subtypes are generated consecutively during chick retinal development from progenitors that are sensitive to follistatin. BMC Dev Biol 8, 1–21. 10.1186/1471-213X-8-46

Field, G.D., Gauthier, J.L., Sher, A., Greschner, M., Machado, T.A., Jepson, L.H., Shlens, J., Gunning, D.E., Mathieson, K., Dabrowski, W., Paninski, L., Litke, A.M., Chichilnisky, E.J., 2010. Functional connectivity in the retina at the resolution of photoreceptors. Nature 467, 673–677. 10.1038/nature09424

Fischer, A.J., Stanke, J.J., Aloisio, G., Hoy, H., Stell, W.K., 2007. Heterogeneity of horizontal cells in the chicken retina. J Comp Neurol 500, 1154–1171. 10.1002/cne.21236

Gallego, A., 1986. Chapter 7 Comparative studies on horizontal cells and a note on microglial cells. Progress in Retinal Research 5, 165–206. 10.1016/0278-4327(86)90010-6

Günther, A., Dedek, K., Haverkamp, S., Irsen, S., Briggman, K.L., Mouritsen, H., 2021. Double Cones and the Diverse Connectivity of Photoreceptors and Bipolar Cells in an Avian Retina. J Neurosci 41, 5015–5028. 10.1523/JNEUROSCI.2495-20.2021

Günther, A., Einwich, A., Sjulstok, E., Feederle, R., Bolte, P., Koch, K.-W., Solov’yov, I.A., Mouritsen, H., 2018. Double-Cone Localization and Seasonal Expression Pattern Suggest a Role in Magnetoreception for European Robin Cryptochrome 4. Curr Biol 28, 211–223.e4. 10.1016/j.cub.2017.12.003

Günther, A., Haverkamp, S., Irsen, S., Watkins, P.V., Dedek, K., Mouritsen, H., Briggman, K.L., 2024. Species-specific circuitry of double cone photoreceptors in two avian retinas. Commun Biol 7, 992. 10.1038/s42003-024-06697-2

Hahn, J., Monavarfeshani, A., Qiao, M., Kao, A.H., Kölsch, Y., Kumar, A., Kunze, V.P., Rasys, A.M., Richardson, R., Wekselblatt, J.B., Baier, H., Lucas, R.J., Li, W., Meister, M., Trachtenberg, J.T., Yan, W., Peng, Y.-R., Sanes, J.R., Shekhar, K., 2023. Evolution of neuronal cell classes and types in the vertebrate retina. Nature 624, 415–424. 10.1038/s41586-023-06638-9

Helmstaedter, M., Briggman, K.L., Turaga, S.C., Jain, V., Seung, H.S., Denk, W., 2013. Connectomic reconstruction of the inner plexiform layer in the mouse retina. Nature 500, 168–174. 10.1038/nature12346

Hore, P.J., Mouritsen, H., 2016. The Radical-Pair Mechanism of Magnetoreception. Annu Rev Biophys 45, 299–344. 10.1146/annurev-biophys-032116-094545

Hsiang, J.-C., Shen, N., Soto, F., Kerschensteiner, D., 2024. Distributed feature representations of natural stimuli across parallel retinal pathways. Nat Commun 15, 1920. 10.1038/s41467-024-46348-y

Kalloniatis, M., Fletcher, E.L., 1993. Immunocytochemical localization of the amino acid neurotransmitters in the chicken retina. Journal of Comparative Neurology 336, 174–193. 10.1002/cne.903360203

Keeley, P.W., Kim, J.J., Lee, S.C.S., Haverkamp, S., Reese, B.E., 2017. Random Spatial Patterning of Cone Bipolar Cell Mosaics in the Mouse Retina. Vis Neurosci 34, E002. 10.1017/S0952523816000183

Kerschensteiner, D., 2022. Feature Detection by Retinal Ganglion Cells. Annu. Rev. Vis. Sci. 8, 135–169. 10.1146/annurev-vision-100419-112009

Kolb, H., Perlman, I., Normann, R.A., 1988. Neural organization of the retina of the turtle *Mauremys caspica:* a light microscope and Golgi study. Vis Neurosci 1, 47–72. 10.1017/S0952523800001012

Leeper, H.F., 1978. Horizontal cells of the turtle retina. II. Analysis of interconnections between photoreceptor cells and horizontal cells by light microscopy. Journal of Comparative Neurology 182, 795–809. 10.1002/cne.901820504

Loew, E.R., Govardovskii, V.I., 2001. Photoreceptors and visual pigments in the red-eared turtle, Trachemys scripta elegans. Vis Neurosci 18, 753–757. 10.1017/s0952523801185081

Macosko, E.Z., Basu, A., Satija, R., Nemesh, J., Shekhar, K., Goldman, M., Tirosh, I., Bialas, A.R., Kamitaki, N., Martersteck, E.M., Trombetta, J.J., Weitz, D.A., Sanes, J.R., Shalek, A.K., Regev, A., McCarroll, S.A., 2015. Highly Parallel Genome-wide Expression Profiling of Individual Cells Using Nanoliter Droplets. Cell 161, 1202–1214. 10.1016/j.cell.2015.05.002

Mariani, A.P., 1987. Neuronal and synaptic organization of the outer plexiform layer of the pigeon retina. Am J Anat 179, 25–39. 10.1002/aja.1001790105

Mouritsen, H., 2018. Long-distance navigation and magnetoreception in migratory animals. Nature 558, 50–59. 10.1038/s41586-018-0176-1

Nemitz, L., Dedek, K., Janssen-Bienhold, U., 2021. Synaptic Remodeling in the Cone Pathway After Early Postnatal Horizontal Cell Ablation. Front Cell Neurosci 15, 657594. 10.3389/fncel.2021.657594

Nemitz, L., Dedek, K., Janssen-Bienhold, U., 2019. Rod Bipolar Cells Require Horizontal Cells for Invagination Into the Terminals of Rod Photoreceptors. Front Cell Neurosci 13, 423. 10.3389/fncel.2019.00423

Peichl, L., Gonzalez-Soriano, J., 1994. Morphological types of horizontal cell in rodent retinae: a comparison of rat, mouse, gerbil, and guinea pig. Vis Neurosci 11, 501–17.

Schindelin, J., Arganda-Carreras, I., Frise, E., Kaynig, V., Longair, M., Pietzsch, T., Preibisch, S., Rueden, C., Saalfeld, S., Schmid, B., Tinevez, J.-Y., White, D.J., Hartenstein, V., Eliceiri, K., Tomancak, P., Cardona, A., 2012. Fiji: an open-source platform for biological-image analysis. Nat Methods 9, 676–682. 10.1038/nmeth.2019

Seifert, M., Baden, T., Osorio, D., 2020. The retinal basis of vision in chicken. Semin Cell Dev Biol 106, 106–115. 10.1016/j.semcdb.2020.03.011

Seifert, M., Roberts, P.A., Kafetzis, G., Osorio, D., Baden, T., 2023. Birds multiplex spectral and temporal visual information via retinal On- and Off-channels. Nat Commun 14, 5308. 10.1038/s41467-023-41032-z

Shekhar, K., Sanes, J.R., 2021. Generating and Using Transcriptomically Based Retinal Cell Atlases. Annu Rev Vis Sci 7, 43–72. 10.1146/annurev-vision-032621-075200

Sonntag, S., Dedek, K., Dorgau, B., Schultz, K., Schmidt, K.-F., Cimiotti, K., Weiler, R., Löwel, S., Willecke, K., Janssen-Bienhold, U., 2012. Ablation of Retinal Horizontal Cells from Adult Mice Leads to Rod Degeneration and Remodeling in the Outer Retina. J. Neurosci. 32, 10713–10724. 10.1523/JNEUROSCI.0442-12.2012

Spinelli, M., Harnecker, A.A., Block, C.T., Lindenthal, L., Schuhmann, F., Greschner, M., Janssen-Bienhold, U., Dedek, K., Puller, C., 2024. The first interneuron of the mouse visual system is tailored to the natural environment through morphology and electrical coupling. iScience 27. 10.1016/j.isci.2024.111276

Ströh, S., Puller, C., Swirski, S., Hölzel, M.-B., van der Linde, L.I.S., Segelken, J., Schultz, K., Block, C., Monyer, H., Willecke, K., Weiler, R., Greschner, M., Janssen-Bienhold, U., Dedek, K., 2018. Eliminating Glutamatergic Input onto Horizontal Cells Changes the Dynamic Range and Receptive Field Organization of Mouse Retinal Ganglion Cells. J Neurosci 38, 2015–2028. 10.1523/JNEUROSCI.0141-17.2018

Sümbül, U., Song, S., McCulloch, K., Becker, M., Lin, B., Sanes, J.R., Masland, R.H., Seung, H.S., 2014. A genetic and computational approach to structurally classify neuronal types. Nat Commun 5, 3512. 10.1038/ncomms4512

Tanabe, K., Takahashi, Y., Sato, Y., Kawakami, K., Takeichi, M., Nakagawa, S., 2006. Cadherin is required for dendritic morphogenesis and synaptic terminal organization of retinal horizontal cells. Development 133, 4085–4096. 10.1242/dev.02566

Tetenborg, S., Yadav, S.C., Hormuzdi, S.G., Monyer, H., Janssen-Bienhold, U., Dedek, K., 2017. Differential Distribution of Retinal Ca2+/Calmodulin-Dependent Kinase II (CaMKII) Isoforms Indicates CaMKII-β and -δ as Specific Elements of Electrical Synapses Made of Connexin36 (Cx36). Front. Mol. Neurosci. 10. 10.3389/fnmol.2017.00425

Thoreson, W.B., Dacey, D.M., 2019. Diverse Cell Types, Circuits, and Mechanisms for Color Vision in the Vertebrate Retina. Physiological Reviews 99, 1527–1573. 10.1152/physrev.00027.2018

Thoreson, W.B., Mangel, S.C., 2012. Lateral interactions in the outer retina. Progress in Retinal and Eye Research 31, 407–441. 10.1016/j.preteyeres.2012.04.003

Tranchina, D., Gordon, J., Shapley, R., 1983. Spatial and temporal properties of luminosity horizontal cells in the turtle retina. Journal of General Physiology 82, 573–598. 10.1085/jgp.82.5.573

Trümpler, J., Dedek, K., Schubert, T., de Sevilla Müller, L.P., Seeliger, M., Humphries, P., Biel, M., Weiler, R., 2008. Rod and cone contributions to horizontal cell light responses in the mouse retina. J. Neurosci. 28, 6818–6825. 10.1523/JNEUROSCI.1564-08.2008

Wässle, H., Puller, C., Müller, F., Haverkamp, S., 2009. Cone contacts, mosaics, and territories of bipolar cells in the mouse retina. J Neurosci 29, 106–117. 10.1523/JNEUROSCI.4442-08.2009

Wiltschko, W., Munro, U., Ford, H., Wiltschko, R., 1993. Red light disrupts magnetic orientation of migratory birds. Nature 364, 525–527. 10.1038/364525a0

Witkovsky, P., 2000. Photoreceptor classes and transmission at the photoreceptor synapse in the retina of the clawed frog, Xenopus laevis. Microscopy Research and Technique 50, 338–346. 10.1002/1097-0029(20000901)50:5<338::AID-JEMT3>3.0.CO;2-I

Xu, J., Jarocha, L.E., Zollitsch, T., Konowalczyk, M., Henbest, K.B., Richert, S., Golesworthy, M.J., Schmidt, J., Déjean, V., Sowood, D.J.C., Bassetto, M., Luo, J., Walton, J.R., Fleming, J., Wei, Y., Pitcher, T.L., Moise, G., Herrmann, M., Yin, H., Wu, H., Bartölke, R., Käsehagen, S.J., Horst, S., Dautaj, G., Murton, P.D.F., Gehrckens, A.S., Chelliah, Y., Takahashi, J.S., Koch, K.-W., Weber, S., Solov’yov, I.A., Xie, C., Mackenzie, S.R., Timmel, C.R., Mouritsen, H., Hore, P.J., 2021. Magnetic sensitivity of cryptochrome 4 from a migratory songbird. Nature 594, 535–540. 10.1038/s41586-021-03618-9

Yamagata, M., Yan, W., Sanes, J.R., 2021. A cell atlas of the chick retina based on single-cell transcriptomics. Elife 10, e63907. 10.7554/eLife.63907

Yoshimatsu, T., Bartel, P., Schröder, C., Janiak, F.K., St-Pierre, F., Berens, P., Baden, T., 2021. Ancestral circuits for vertebrate color vision emerge at the first retinal synapse. Science Advances 7, eabj6815. 10.1126/sciadv.abj6815

Zapka, M., Heyers, D., Hein, C.M., Engels, S., Schneider, N.-L., Hans, J., Weiler, S., Dreyer, D., Kishkinev, D., Wild, J.M., Mouritsen, H., 2009. Visual but not trigeminal mediation of magnetic compass information in a migratory bird. Nature 461, 1274–1277. 10.1038/nature08528

